# MicroED structure of the human adenosine receptor determined from a single nanocrystal in LCP

**DOI:** 10.1101/2020.09.27.316109

**Authors:** Michael W. Martynowycz, Anna Shiriaeva, Xuanrui Ge, Johan Hattne, Brent L. Nannenga, Vadim Cherezov, Tamir Gonen

**Affiliations:** Howard Hughes Medical Institute, University of California Los Angeles, Los Angeles, CA, USA; Department of Biological Chemistry, University of California Los Angeles, Los Angeles, CA, USA; Bridge Institute, USC Michelson Center for Convergent Biosciences, University of Southern California, Los Angeles, CA, USA; Department of Chemistry, University of Southern California, Los Angeles, CA, USA; Mork Family Department of Chemical Engineering and Materials Science, University of Southern California, Los Angeles, CA, USA; Chemical Engineering, School for Engineering of Matter, Transport and Energy, Arizona State University, 501 E. Tyler Mall, PO Box 876106, Tempe, AZ 85287, United States; Biodesign Center for Applied Structural Discovery, Biodesign Institute, Arizona State University, 727 East Tyler Street, Tempe, AZ 85287, United States; Department of Physiology, University of California Los Angeles, Los Angeles, CA, USA

**Author notes:** Corresponding authors. (V.C.); (T.G.).

## Abstract

G Protein-Coupled Receptors (GPCRs), or 7-transmembrane receptors, are a superfamily of membrane proteins that are critically important to physiological processes in the human body. Determining high-resolution structures of GPCRs without signaling partners bound requires crystallization in lipidic cubic phase (LCP). GPCR crystals grown in LCP are often too small for traditional X-ray crystallography. These microcrystals are ideal for investigation by microcrystal electron diffraction (MicroED), but the gel-like nature of LCP makes traditional approaches to MicroED sample preparation insurmountable. Here we show that the structure of a human A_2A_ adenosine receptor can be determined by MicroED after converting the LCP into the sponge phase followed by cryoFIB milling. We determined the structure of the A_2A_ receptor to 2.8 Å resolution and resolved an antagonist in its orthosteric ligand-binding site as well as 4 cholesterol molecules bound to the receptor. This study lays the groundwork for future GPCR structural studies using single microcrystals that would otherwise be impossible by other crystallographic methods.

**One sentence summary:** FIB milled LCP-GPCR structure determined by MicroED

## Main text

G protein-coupled receptors (GPCRs) constitute the largest and highly diverse membrane protein superfamily in the human genome represented by over 800 members (*1, 2*). Expressed on the cell surface plasma membrane, receptors function as cellular gatekeepers, transmitting signals inside the cell in response to a variety of signaling molecules and environmental cues. GPCR-mediated signaling pathways play a key role in all vital physiological systems as well as pathophysiological conditions including cancer, cardiovascular diseases, immune and metabolic disorders, pain and addiction, and others (*3*). Because of their fundamental roles in health and disease, GPCRs have been recognized as important drug targets, with over 30% of all approved therapeutic drugs acting via these receptors (*4*).

Adenosine A_2A_ receptor (A_2A_AR) is a prototypical and one of the most extensively studied GPCR (*5*). It expresses broadly in the central nervous system and peripheral tissues and responds to an extracellular neuromodulator adenosine, mediating a range of physiological processes including sleep regulation, angiogenesis, and immunosuppression. A_2A_AR agonists are used clinically in pharmacological stress testing because of their vasodilatory effects (*5*), while antagonists have been considered as potential candidates for the treatment of Parkinson’s disease and other neurodegenerative disorders (*6*), as well as, more recently, as promising agents for cancer immunotherapy (*7*).

Structure-based drug discovery (SBDD) and optimization require accurate atomic models (*8, 9*). Recent advances in single particle analysis (SPA) in an electron cryo-microscopy (cryoEM) have allowed for high-resolution structures of several GPCRs to be determined, most of which have been obtained in complex with the G-protein because of the size limitations in SPA (*10, 11*). Nevertheless, crystallography remains the only approach of studying the receptors in their inactive state, without a signaling partner bound, in complex with antagonists or inverse agonists. The majority of GPCR structures have been determined using crystallization in a lipidic cubic phase (LCP) (*12, 13*).

Determining GPCR structures using traditional X-ray crystallography is challenging. Extracting crystals from the viscous LCP is difficult, and many membrane protein crystals only grow to a few micrometers in their largest dimension. Even when relatively large crystals are available, soaking drugs into them is not always feasible, limiting SBDD applications for GPCRs (*14*). Therefore, structural investigations of GPCRs were greatly facilitated by the advent of X–ray free electron lasers (XFEL) with injector-based LCP delivery systems (*10, 15, 16*). These approaches took many years to develop, XFEL sources are costly, and access to XFEL beamtime is highly competitive. Furthermore, millions of individual crystals are typically used during an XFEL experiment, data from a few thousand are then merged to generate a structure. For these reasons we explored the use of a relatively new MicroED method for determining the structure of a GPCR from crystals grown in LCP.

Microcrystal electron diffraction (MicroED) is ideally suited to study protein nanocrystals of soluble proteins (*17*–*19*), membrane proteins (*20*–*22*) and small molecules (*19, 23*–*25*). However, recent investigations attempting to extract crystals from the viscous lipid-matrix have demonstrated that traditional sample preparation methods are ill-equipped to isolate well-diffracting crystals for MicroED experiments (*26, 27*). Clearly, further important improvements were required to make GPCR crystals grown in LCP amenable to MicroED analyses.

Here, we combine cryogenic focused ion-beam (FIB) milling with MicroED to determine the structure of the GPCR A_2A_ adenosine receptor (A_2A_AR) embedded in LCP from a single microcrystal. To facilitate crystallization, the A_2A_AR was fused with apocytochrome b_562_RIL into its third intracellular loop, and its C-terminal residues 317-412 were truncated (A_2A_AR-BRIL-ΔC, hereafter A_2A_AR) (*28*). In order to obtain a thin layer of LCP containing GPCR crystals onto a TEM grid, the gel-phased LCP was converted to the sponge phase by mixing the LCP with a sponge phase-inducing agent (*29*). Only after LCP conversion into the sponge phase, blotting away the excess solvent, and FIB milling crystals into thin lamellae, were we able to collect continuous rotation MicroED data and determine the high-resolution structure.

Several methods for grid preparation were attempted before well-diffracting crystals were obtained. In a recent report we demonstrated that mammalian voltage dependent anion channel (mVDAC) crystals grown in the presence of lipid bicelles were amenable to FIB milling and subsequent structure determination by MicroED when crystals were transferred onto a TEM grid at high humidity (*22*). Although we attempted direct crystal transfer for A_2A_AR, that strategy did not yield satisfactory results. This is likely because an LCP is more viscous and difficult to spread out in a thin layer compared with a bicellar preparation. With extensive trial and error we succeeded in transferring crystals onto grids but at the expense of crystal dehydration leading to poor diffraction (Figures 1A, S1, S2).

**Figure 1.**
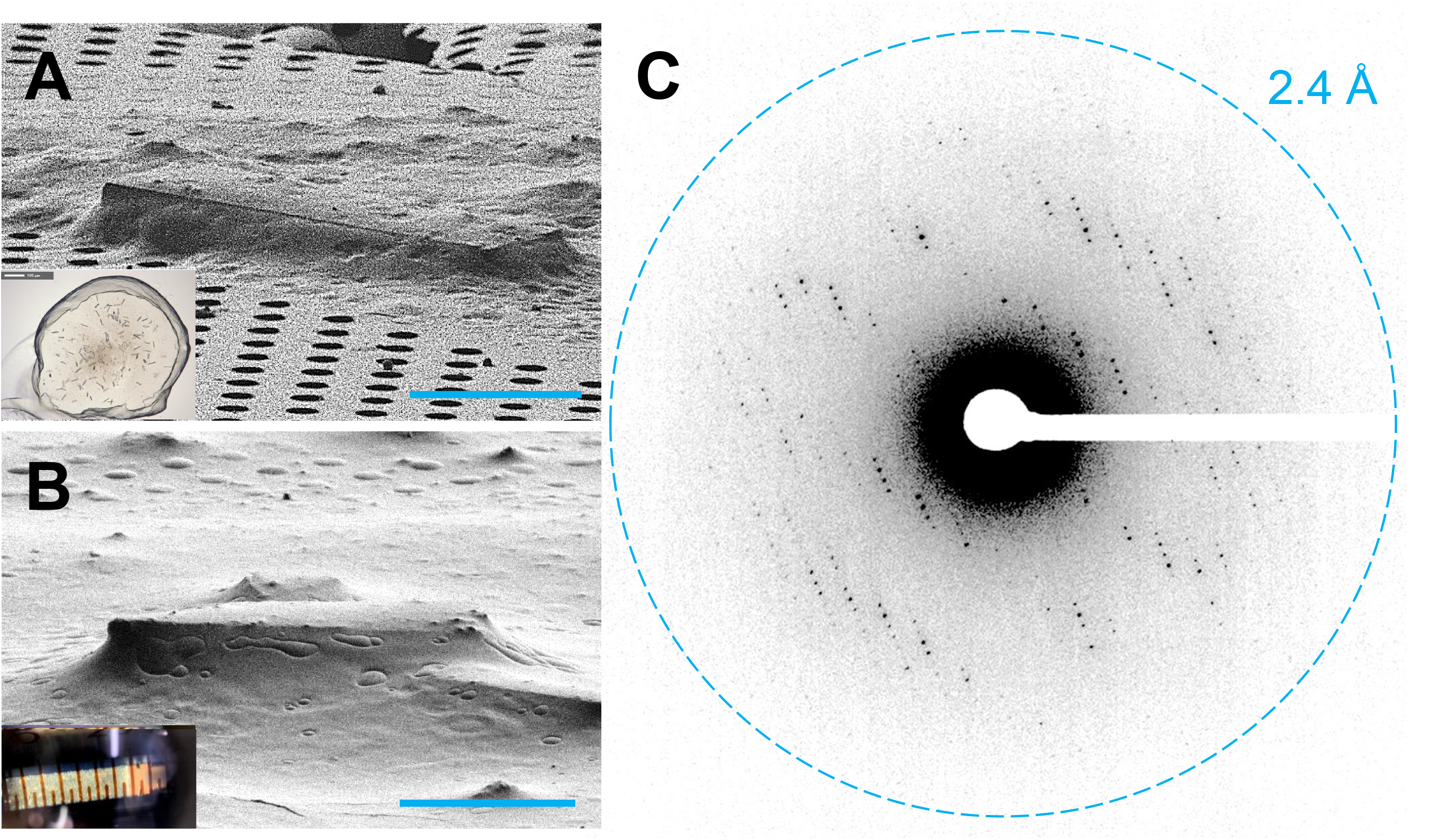
Preparing GPCR crystals for MicroED data collection. (A) An A_2A_AR crystal looped from a glass sandwich plate and placed on a TEM grid viewed in the FIB at 18°. Typical crystallization drop used for looping GPCR crystals shown inset. (B) FIB image of an A_2A_AR crystal from a syringe of microcrystals in the sponge phase. Typical syringe of sponge-phase A_2A_AR crystals shown as inset. (C) MicroED data taken from an A_2A_AR crystal lamella, showing clear spots out to 2.7 Å at 0°. A continuous rotation MicroED dataset was taken from this crystal. Scale bars in (A) and (B) correspond to 10 µm.

Conversion of the LCP phase from the gel to the sponge phase by application of sponge-inducing agents or treatment with a lipase was explored. We recently described this method and demonstrated that some diffraction data could be obtained from a GPCRcrystal (*26*). In that report, aliquots of GPCR crystals grown in LCP were applied to a TEM grid and covered by a lipase solution, MPD, or PEG on top of the LCP bolus. Conversion of the LCP to the sponge phase was completed in about 1 hour. The grid was then observed in the transmission electron microscope, revealing large areas that were too thick for the electron beam to penetrate. Small crystalline wedges sticking out of the thick LCP matrix were identified. These crystalline wedges appeared dehydrated and only diffracted to around 4-5 Å resolution and lasted for just a few exposures before they were destroyed by radiation damage. We surmised that the long incubation time for conversion of the LCP from the gel to the sponge phase may have damaged the fragile crystals so further modification to this method had to be developed.

The procedure for conversion of the LCP phase from the gel to the sponge phase was modified as follows. Microcrystals of A_2A_AR-BRIL were grown in a syringe as described (*30, 31*). The precipitant solution was ejected from the syringe, leaving behind only the LCP with embedded microcrystals. Small portions of PEG400 100% were added to the LCP stepwise (∼5% of the total LCP volume per step), and mixed mechanically back and forth between two syringes until a homogeneous mixture was achieved (Figure 1B, inset, Figure S1). This continued until the gel-like LCP was converted into a liquid-like sponge phase and the syringe could dispense the mixture through a narrow needle without additional force being necessary. Microcrystals remained visually intact throughout this procedure. A very small portion (∼0.5 μL) of the resultant sponge phase was applied to a TEM grid inside of a vitrification robot at room temperature and 90% humidity. The grid was blotted from the front and then the back, plunged into liquid ethane, and stored under cryogenic conditions.

Vitrified grids with the sponge-A_2A_AR microcrystals were loaded into a dual-beam focused ion-beam, scanning electron microscope (FIB/SEM) (*32, 33*). To our satisfaction, many small crystals were visible on the grid using low-magnification imaging in the FIB/SEM (Figure 1B, Figure S3). Unlike previous preparations, these crystals were well embedded in a homogeneous layer of a sponge phase (Figure 1B versus 1A). The crystals were milled using a focused ion beam of gallium into a thin lamellae approximately 200 nm thick (Figures S2, S3). The grids were loaded into a Thermo-Fisher Titan Krios, where the lamellae prepared in the FIB/SEM were identified using low-magnification montaging. A single diffraction pattern without tilting or rotation was acquired from each lamella to assess their diffraction quality, and we were happy to find that all prepared lamella diffracted. Four of six lamellae diffracted better than 5 Å. Out of these four, one lamellae diffracted to ∼4.5 Å, two to 3.5 Å, and one to better than 3 Å (Figures 1C, S3). We collected continuous rotation MicroED (*18*) data from the best four lamellae over wedges of approximately 70° (Movie S1).

The MicroED structure of A_2A_AR was determined from a single nanocrystal. The total diffracting volume was less than 1 μm^3^ (0.2×2×2 μm) making this ∼2-3 orders of magnitude smaller than any GPCR crystals used for structure determination to date. The integrated data from the best lamella resulted in a dataset to 2.8 Å resolution, an overall I/sI of 7.1, and a completeness of approximately 80% through the resolution shells. This dataset was indexed in space group C 2 2 2_1_ (#20) with unit cell dimensions of (a, b, c) = 40.0 Å, 180.5 Å, 139.7 Å, and angles (α, β, Ɣ) = 90° consistent with prior reports (*28, 30*). Molecular replacement was conducted using the PDB 4EIY (*28*) as a search model after removing all ligands and converting all side chains into alanines, resulting in a single, unambiguous solution. Initial maps following molecular replacement showed clear difference peaks where the ZM241385 ligand and other lipids were identified in other structures of this protein (*28, 30*). The molecular replacement solution was refined using electron scattering factors until convergence and the density was inspected (Figure 2 and S4-8). One ZM241385 ligand (ZMA) and four cholesterols could be accurately placed in the difference maps (Figure S6, Movie S2-5).

**Figure 2.**
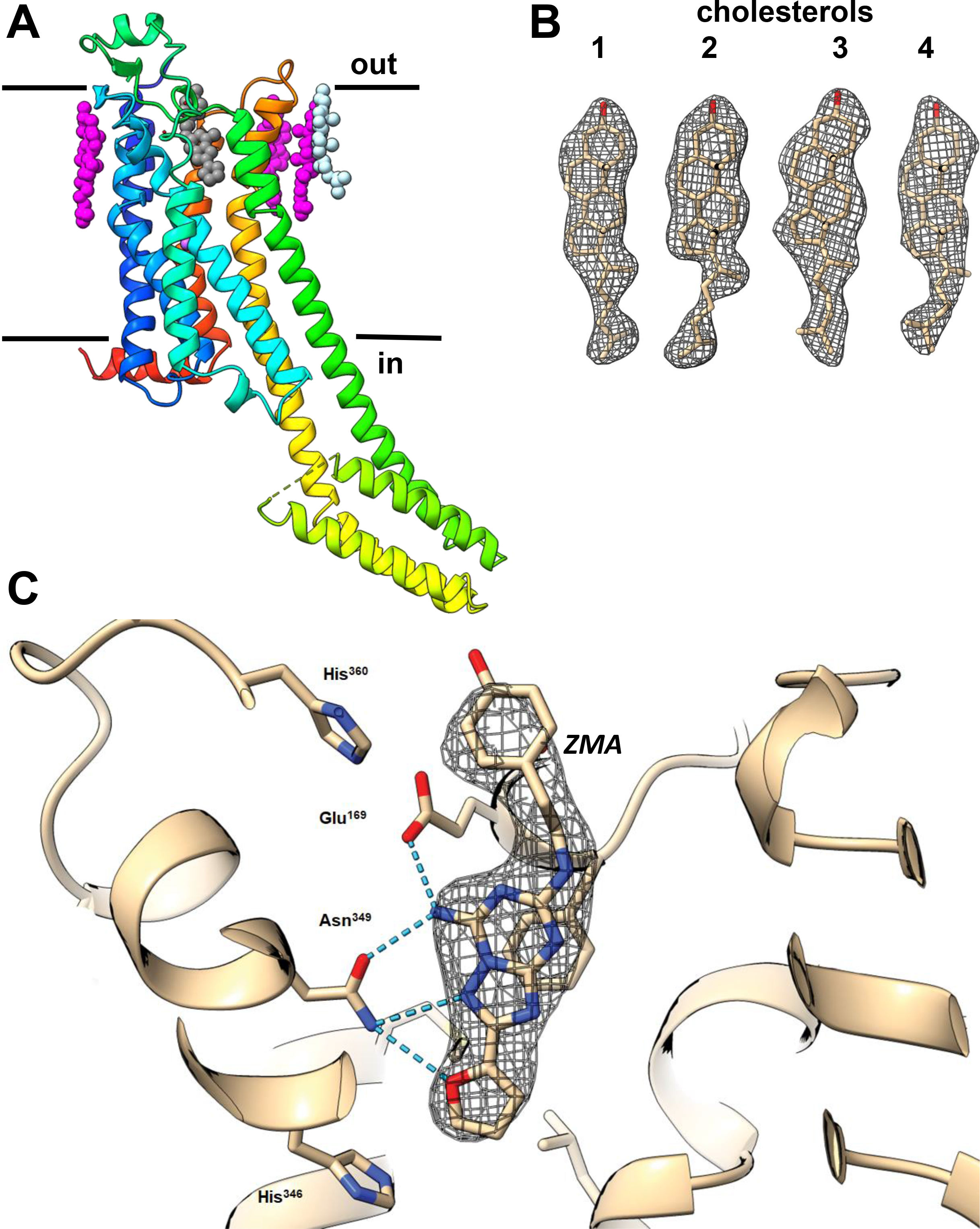
The structure of A_2A_AR by MicroED. (A) Cartoon representation of the A_2A_AR GPCR structure depicted in rainbow viewed from the side. The membrane bilayer as indicated. Ligands are shown in sphere representation, where the ZM241385 antagonist is in black, known cholesterol positions in magenta, and the newly determined cholesterol position in light blue. (B) Density for the four cholesterol molecules (C) Magnified binding pocket of the ZM241385 antagonist from the region indicated by a circle in (A). Density is shown around the ligand along with coordination with the protein residues. 2mF_o_-DF_c_ map is contoured at the 1s level.

The overall structure of A_2A_AR is consistent with past reports (Figures 2, S4) (28, 30). The protein folds into a 7TM topology with the N terminus located on the extracellular side of the membrane and an intracellular helix 8 at the C terminus running parallel to the lipid membrane. The density for all receptor loops and all four disulfide bonds are well defined. TMs 5 and 6 are elongated, protruding well beyond the surface of the cytoplasmic lipid layer where they normally engage the G-protein (*34*). On the extracellular side, four cholesterol molecules were observed per monomer. The antagonist ZM241385 was observed in the orthosteric ligand-binding site inside the 7TM bundle (Figures S5, S6).

The 2.8 Å resolution structure was refined to an R_work_/R_free_ of 24.8/28.8%, and good overall geometry (Table 1). Interestingly, we identified one additional cholesterol in this structure, interacting with Phe183, Phe258 and two other cholesterols, when compared to the previously determined structure of A_2A_AR (PDB 4EIY). Here, the site was occupied by an oleic acid rather than a cholesterol (Figure 2A, light blue and 2B) (*28*). The additional cholesterol identified forms a cluster with two other cholesterols that mediate crystal contacts between two receptor monomers (Movie S4). The location of the bound antagonist, ZM241385 (ZMA), and coordination to the protein was accurately determined and verified in the omit maps (Figures 2C, S4-6).

**Table 1.**
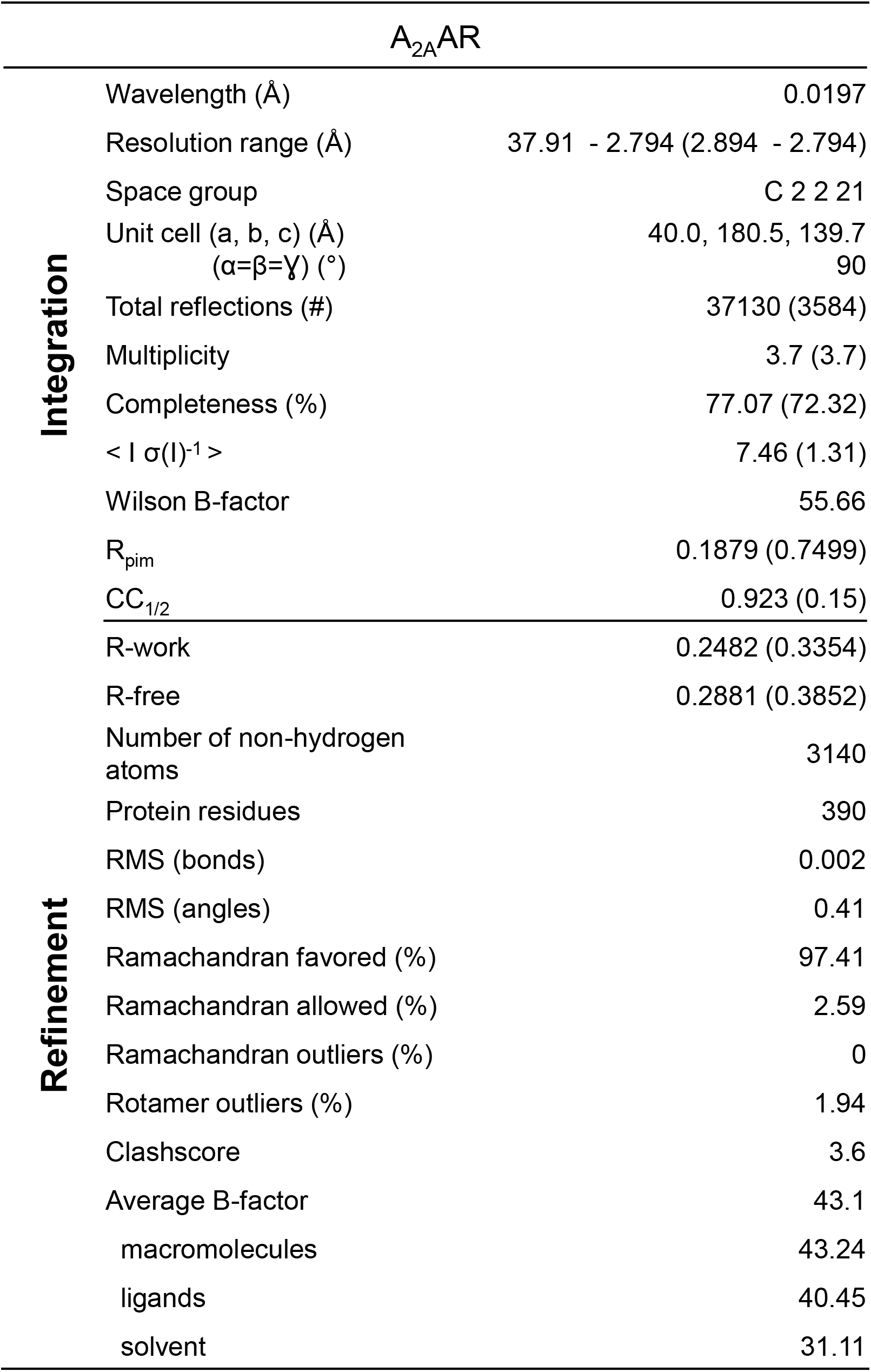
MicroED crystallographic table for A_2A_AR.

MicroED data was collected using an extremely low exposure to minimize radiation damage (*35*). Recent quantification of radiation damage in MicroED studied the effect of exposure of a crystalline sample to the electron beam. The most sensitive to site-specific damage were disulfide bridges, which showed signs of deterioration even at an average exposure of ∼3 e^-^Å^-2^. The MicroED structure of A_2A_AR was determined using an average exposure of only 2 e^-^Å^-2^ (∼7.4 MGy) and as expected the density for all four disulfide bridges (C71-C159; C74-C146; C77-C166; C355-C358) remained intact (Figure S8). This observation suggests that FIB milling did not inflict significant additional radiation damage on the sample.

Because of their rod-like shape, these A_2A_AR crystals adopt a preferred orientation on the EM grids. Thorough attempts were made to merge the additional datasets collected to increase completeness in the lower resolution shells of our dataset. Doing so only increased completeness to 84% at 3.8 Å resolution, resulting in large degradation of all the crystallographic statistics likely because of variations in unit cell parameters. Prior studies have shown that crystals with preferred orientations on the EM grids will suffer from a systematic missing wedge of data, for which merging data from additional crystals cannot compensate (*36, 37*). Despite the preferred orientation, a single A_2A_AR lamella yielded 80% completeness and a high quality density map and structure to 2.8 Å resolution.

The structure of A_2A_AR was previously determined at better than 2 Å resolution using XFEL and synchrotron radiation (28, 30). In the case of XFEL, a dataset was assembled from 72,735 single orientation snapshots from crystals with an average volume of 50 μm^3^ per crystal (30), while in the case of synchrotron radiation data were collected from 55 crystals with an average volume of 1,800 μm^3^ per crystal (28). The MicroED structure presented here accomplished by using a single crystal with less than 1 µm^3^ volume which therefore represents a significant improvement in both the number and size of crystallites needed for structure determination of GPCRs and identification of ligands.

We determined the first MicroED structure of a GPCR grown in LCP. This structure was determined from a single crystal to 2.8 Å resolution. The crystals were made amenable to MicroED investigation by converting the LCP into the sponge phase and subsequent FIB milling of the microcrystals spread on a TEM grid in a humidity controlled environment. The structure was derived from a single microcrystal milled to a thin lamellae only 200 nm thick. The total illuminated crystalline volume for a complete, near-atomic dataset was less than 1 µm^3^, a feat essentially impossible using any other crystallographic method. The fully refined structure clearly identified most of the side chains, the bound ligand and four cholesterols even though they were not included in the search model for molecular replacement, testifying to the high quality of the data obtained. Consistent with the above postulate, we did not observe significant signs of site-specific damage as all four disulfide bridges appeared intact in the A_2A_AR density map. The use of sponge phase with microcrystals lays the groundwork for future inquiries of GPCR microcrystals by MicroED. Of particular interest is the availability of soaking experiments conducted on-grid for structure based drug discovery (*38*). This investigation represents a leap forward in the use of MicroED and FIB milling of GPCR microcrystals, and has broad implications for how the structures of membrane proteins grown in LCP may be in solved in the future using even smaller crystals.

## Acknowledgements

This study was supported by the National Institutes of Health P41GM136508 to TG, R35 GM127086 to VC, R01GM124152 to BLN. The Gonen lab is supported by funds from the Howard Hughes Medical Institute.

## Author contributions

MWM, VC and BLN conducted MicroED experiments. A.S. and X.G. purified and crystallized the receptor. Structure determination was conducted by MWN, VC and JH. Figures were prepared by MWM and TG. The paper was written by MWM, VC, and TG with input and contributions from all authors. The project was conceived by VC and TG.

## Data and material availability

Materials and reagents will be made available upon reasonable request to the lead contact. The structure factors and coordinates for the A_2A_AR structure will be deposited in the PDB, and the associated maps will be deposited at the EMDB.

## Supplementary materials

### Materials and methods

#### Protein production

Expression and purification of A_2A_AR, containing BRIL fusion protein in the third intracellular loop and a C-terminal truncation of residues 317-412 (A_2A_AR-BRIL-ΔC), were done as previously described (*30*).

#### Crystallization

Purified and concentrated to 30 mg/mL protein samples of A_2A_AR-BRIL-ΔC in complex with ZM241385 were reconstituted into LCP by mixing with molten lipid using a syringe mixer (*39*). The protein-LCP mixture contained 40% (w/w) protein solution, 54% (w/w) monoolein (Sigma), and 6% (w/w) cholesterol (Sigma). Crystals for MicroED data collection were obtained in 96-well glass sandwich plates (Marienfeld) and in Hamilton gas-tight syringes similarly to (*30, 31, 39*). Precipitant solution contained 50-75 mM sodium thiocyanate, 100 mM sodium citrate pH 4.8, 28% (v/v) PEG 400, 2% (v/v) 2,5-hexanediol. Crystals appeared in 24 hours and reached maximum size of 30-50 μm in plates and 10-20 μm in syringes within 5 days.

#### Grid preparation for MicroED

All experiments used Quantifoil Cu200 R2/2 holey carbon grids. All grids were glow discharged immediately before use. Samples were initially made using crystals grown in glass sandwich plates as described (*39*). After cracking open the glass, a portion of the LCP bolus containing a large group of crystals was picked up by a 100 μm MiTeGen dual thickness micromount and then carefully transferred to a glow discharged grid. The transfer was done by gently sliding the loop along the surface of the grid in an attempt to spread out LCP without breaking the carbon foil. This process was modified to include a humidifier in order to keep the crystals hydrated during the looping process, but ultimately resulted in grids with ice too thick to identify crystals or grid bars. Samples of A_2A_AR-BRIL-ΔC crystals grown in syringes were used to convert LCP into a sponge phase. For this purpose, precipitant solutions were carefully removed from three syringes and the remaining LCP with embedded microcrystals was consolidated together into one syringe (∼20 μL of LCP). Approximately 1 μL of 100% PEG 400 was added into a clean syringe and mixed with the LCP sample by moving it back and forth between two syringes through a coupler containing a gauge 22 needle until homogeneity. The process of adding PEG 400 was repeated several times until the gel-like LCP is converted into a liquid-like sponge phase that could be ejected from a gauge 22s needle without applying any extensive force. Approximately 0.5-1 μL of this sponge phase with microcrystals was placed on a TEM grid inside of a Leica GP2 cryo plunger. The sample chamber was held at 20°C and 90% humidity. The grid was gently blotted from the front for 5 s, the back for 5 s, and then vitrified by plunging into liquid ethane. Grids were stored in liquid nitrogen until use.

#### FIB milling and SEM imaging

All FIB SEM experiments were conducted on a Thermo Fisher Aquilos dual beam FIB SEM as described (*32, 38, 40*). A thick layer of sputter coated platinum approximately 500 nm thick was deposited on the grids in order to protect the samples from the damaging gallium beam. Individual grids were screened using a single low magnification image in the SEM. If the grid was not destroyed and the TEM grid bars were visible through the LCP media, an all-grid map was taken in the MAPS software. Lamellae sites were identified in the SEM map and crystals were verified by imaging at grazing incidence in the FIB. Select crystals were milled as described (*22, 32, 33, 38, 40, 41*). Briefly, crystals were adjusted to eucentric height. Trenches were milled away from the crystals at an angle of 35° to determine the thickness of the underlying LCP material. After trenching, areas of the crystal were removed from either the top or bottom using the ion beam. As the crystal was thinned, the current of the ion-beam was lowered. When the lamella thickness was approximately 300 nm, the lamellae was slowly polished using a 10 pA current until a final thickness of approximately 200 nm was achieved.

#### MicroED data collection, structure determination, and refinement

Continuous rotation MicroED data were collected as described (*18, 35, 42*). Grids with milled crystals were transferred from the Thermo-Fisher Aquilos dual beam FIB/SEM and into a Thermo-Fisher Titan Krios TEM. The Krios was cryogenically cooled to liquid nitrogen temperatures and operated at an accelerating voltage of 300 kV, corresponding to an electron wavelength of 0.0197 Å. All MicroED data were collected on a Thermo-Fisher CetaD 16M CMOS detector. Lamellae were identified in low-magnification montages, where they were apparent as white stripes against an otherwise black background. Identified lamellae were brought to eucentric height and screened for diffraction by taking a 1 s exposure of the lamella in diffraction mode prior to a full data collection. Suitably well diffracting lamellae were isolated using a selected area aperture measuring approximately 2 µm in diameter to reduce background noise from the surrounding areas. MicroED data were collected using an exposure rate of approximately 0.01 e^-^ Å^-2^ s^-1^. MicroED data were collected in wedges of 0.6°, where the detector was readout every 3 s while the stage was rotating at 0.2° s^-1^. Data were collected as MRC stacks of images, and converted to SMV format as described (*32, 43*). The data were indexed, integrated, scaled, and merged in DIALS (*44, 45*).

The structure was determined by molecular replacement in PHASER (*46*) using the PDB 4EIY (*28*) as a search model after removing all ligands and converting all side chains into alanines. The structure was refined iteratively using Phenix (*47*) using electron scattering factors and visual inspection in COOT (*48*) until convergence.

#### Figures and graphic creation

Figures were arranged in Microsoft PowerPoint. Structural models and maps were created using ChimeraX (*49*).

### Supplementary figure and table legends

**Figure S1.**
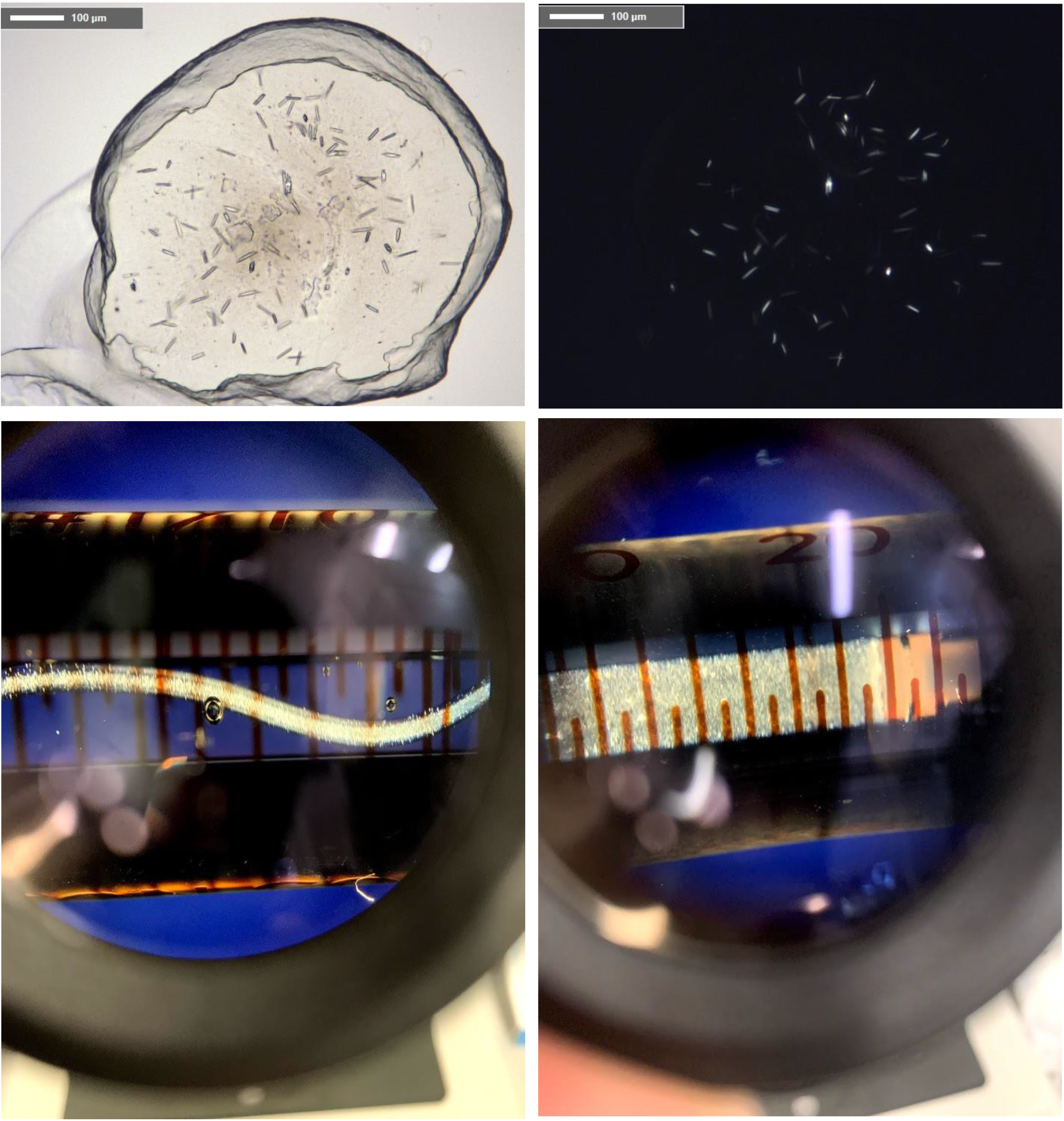
Typical GPCR crystals grown in 96-well glass sandwich plates, syringes, and the GPCR crystals in a syringe after conversion to the sponge phase.

**Figure S2.**
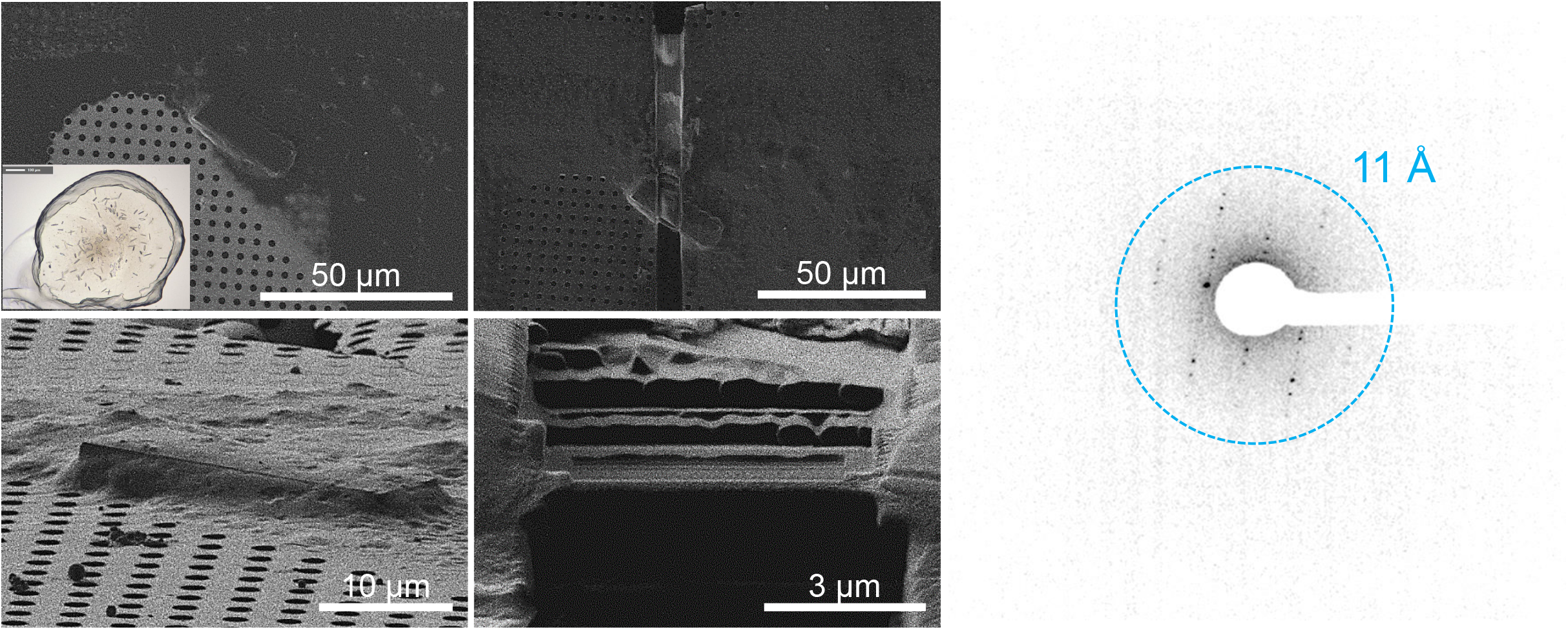
GPCR MicroED data from crystals directly transferred from plates.

**Figure S3.**
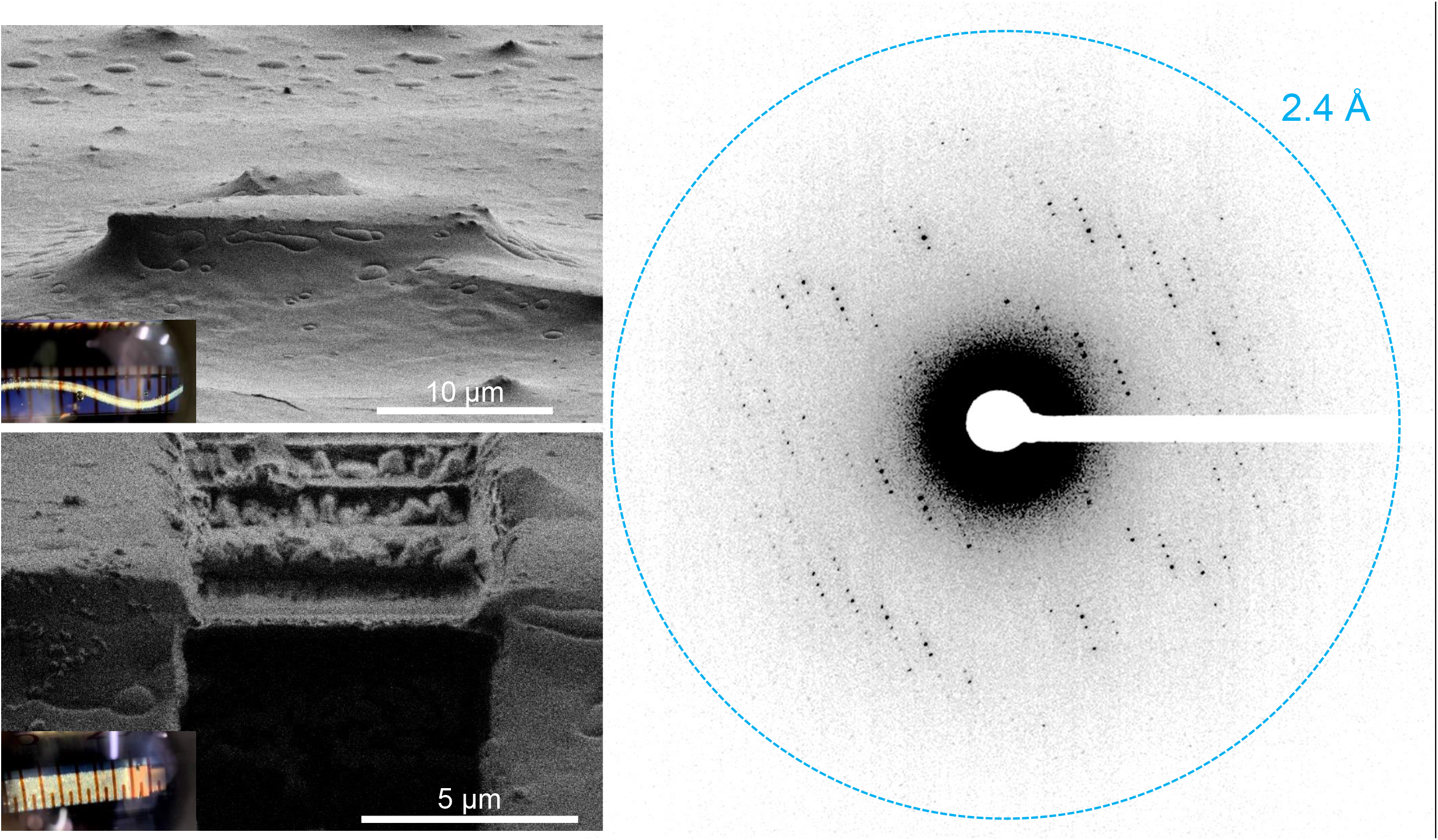
GPCR MicroED data from crystals transferred on the grid in high humidity after conversion of LCP into sponge phase.

**Figure S4.**
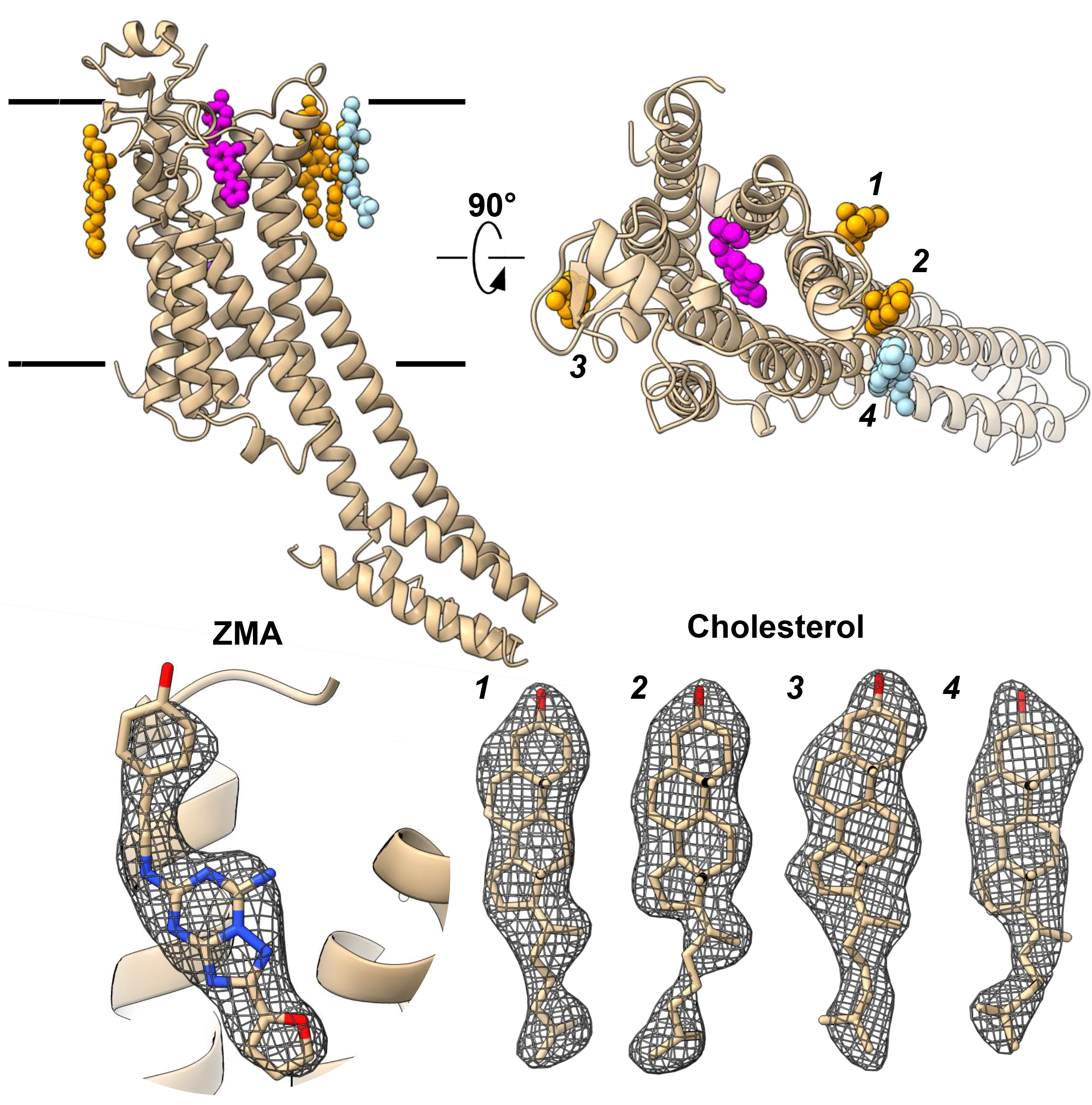
The overall structural architecture of the solved A_2A_AR structure.

**Figure S5.**
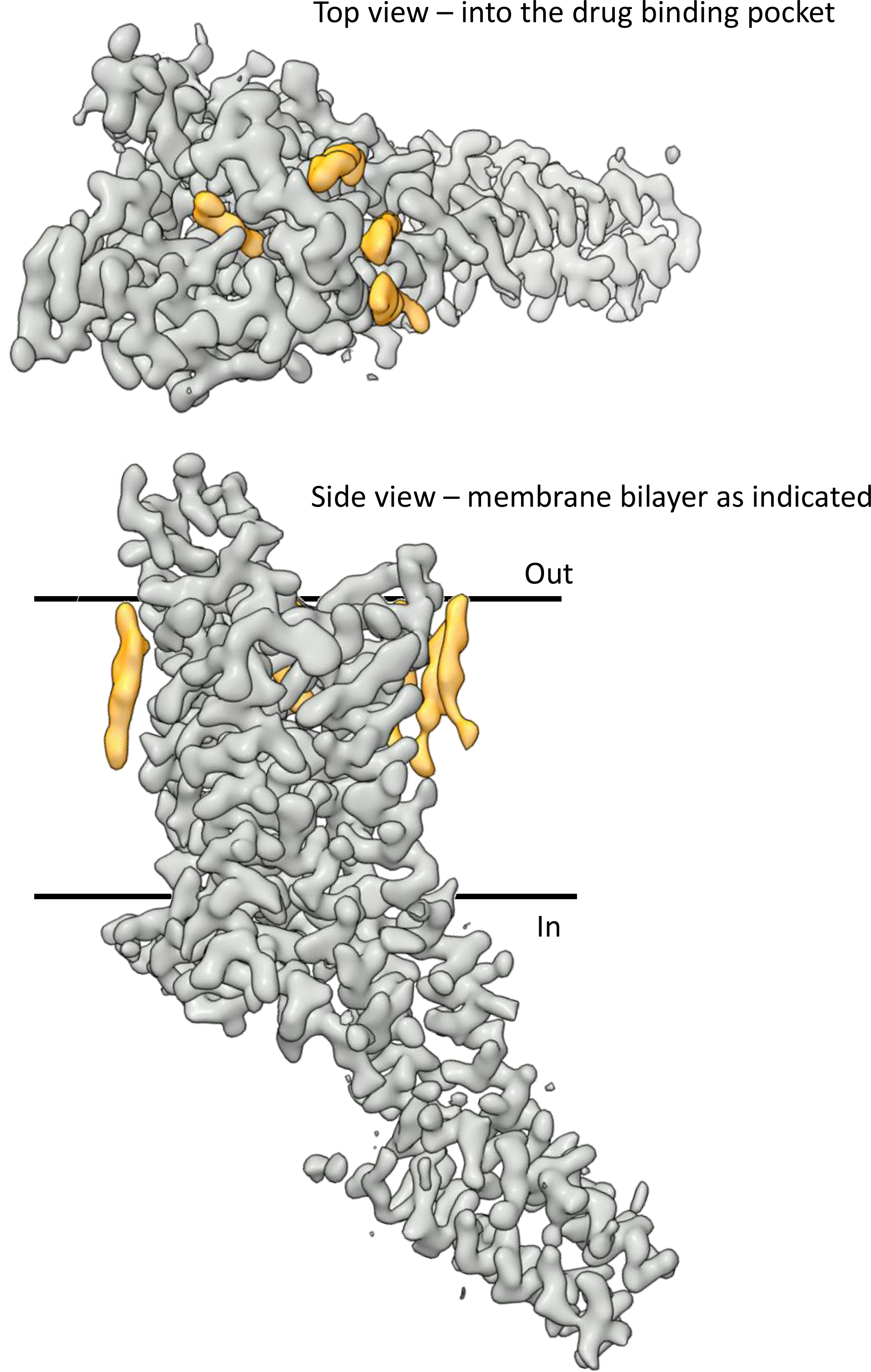
Overall density of the protein (grey) and ligands (yellow) as determined by MicroED.

**Figure S6.**
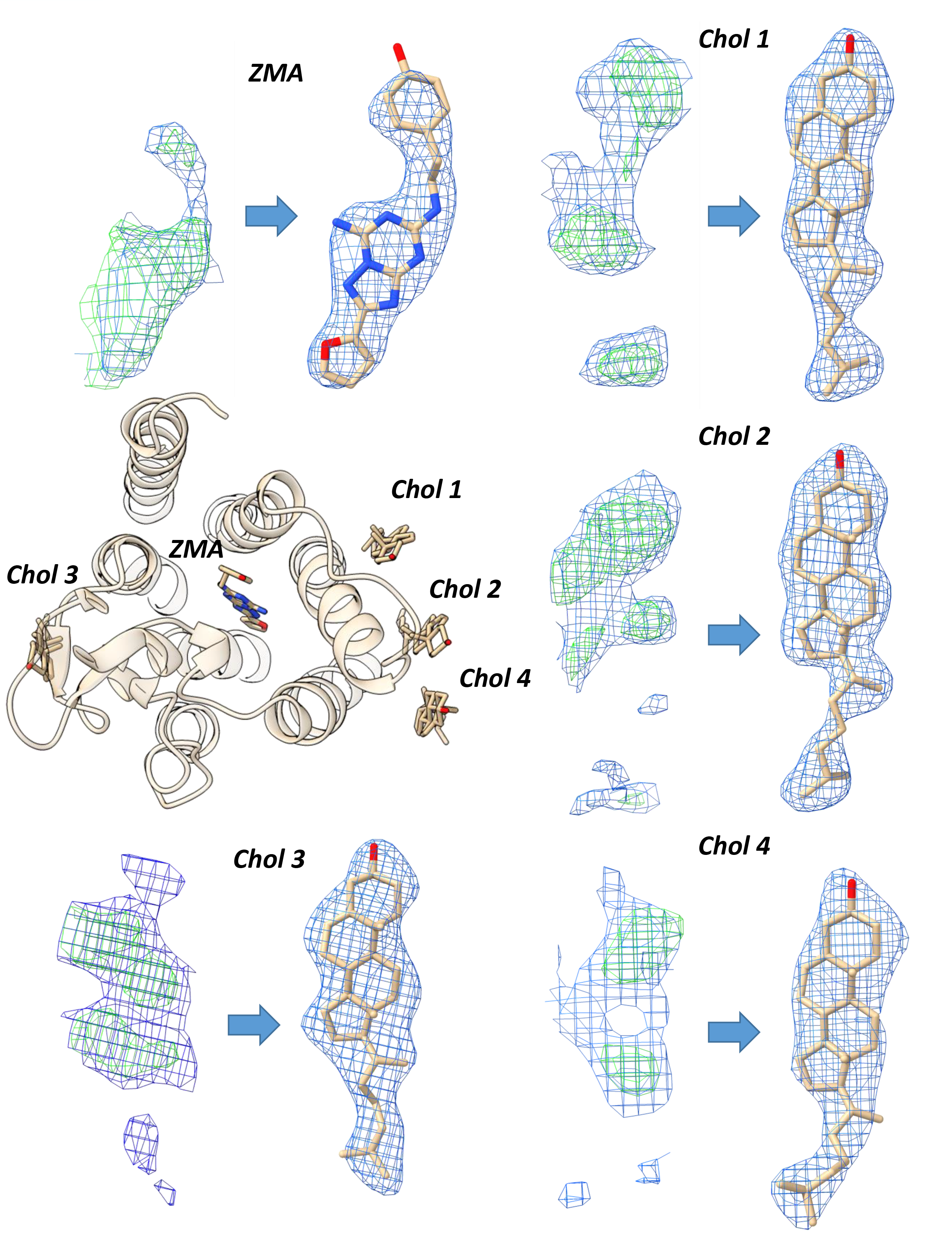
Identification and refinement of ligands not included in the molecular replacement model in the MicroED density. 2mFo-DFc density (grey) is contoured at 0.7s and mFo-DFc is contoured at 2.5s.

**Figure S7.**
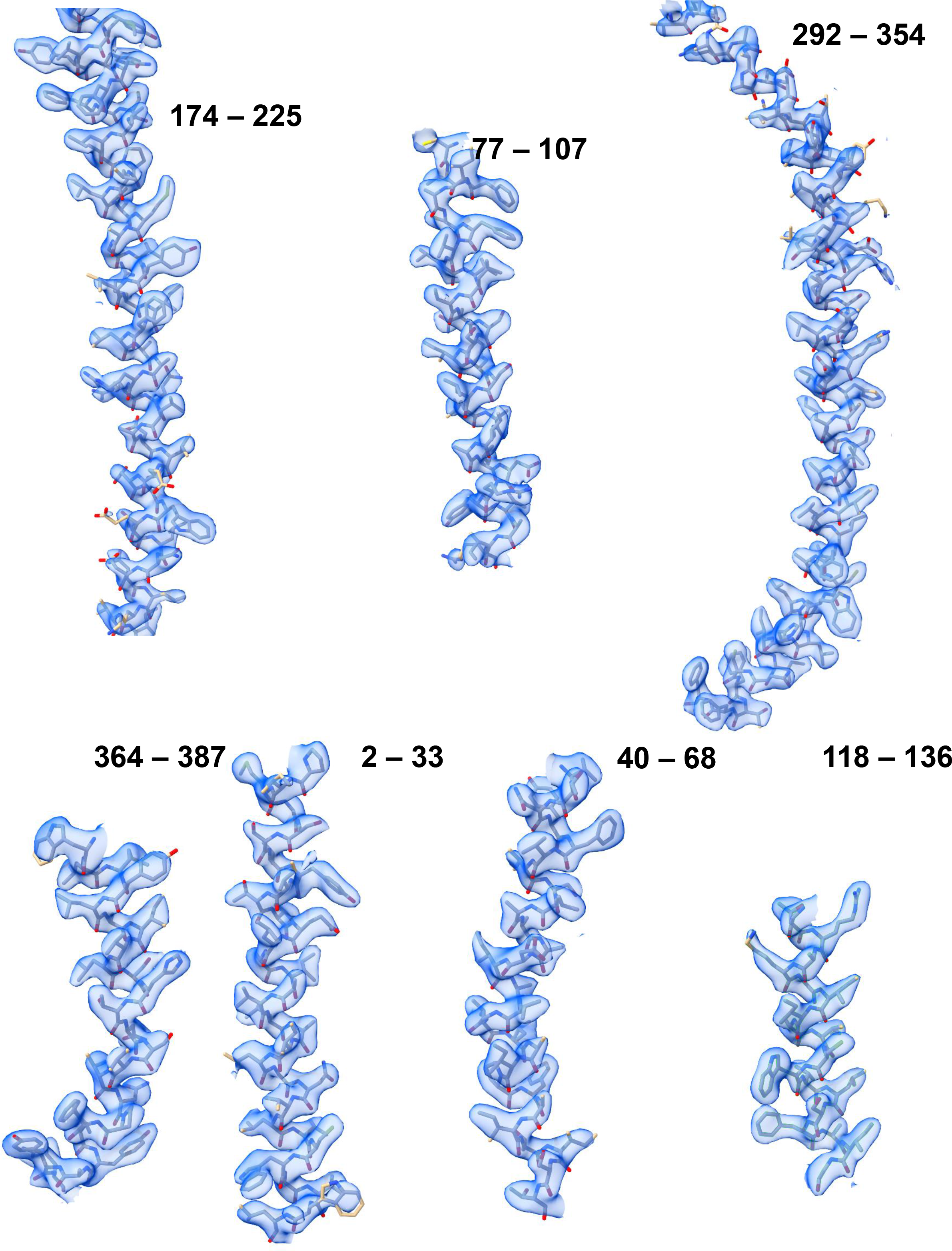
2mFo-DFc Density of the TM helices determined by MicroED, contoured at 1s.

**Figure S8.**
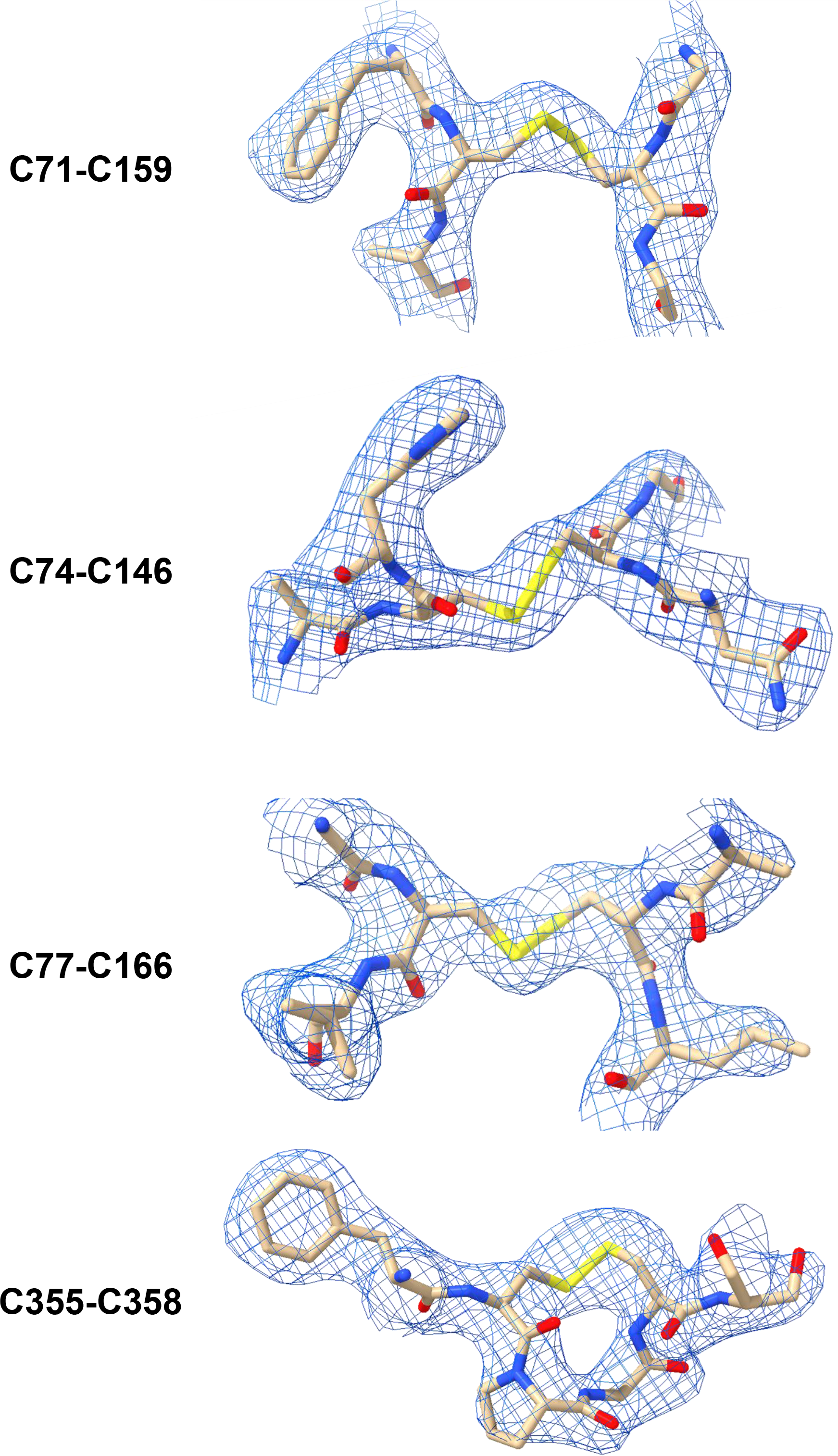
Intact disulfide bridges in the MicroED structure of A_2A_AR. The four disulfide bridges are C71-C159; C74-C146; C77-C166 and C355-C358.

Movie S1. High resolution MicroED data collected from a single GPCR microcrystal. Movie S2. Determined structure of A_2A_AR with associated density.

Movie S3. Structure of A_2A_AR with highlighted ligand locations. Movie S4. Crystal packing of the A_2A_AR structure.

Movie S5. Density around individual residues in the A_2A_AR structure determined by MicroED.

